# The layered costs and benefits of translational redundancy

**DOI:** 10.1101/2022.06.16.495782

**Authors:** Parth K Raval, Wing Yui Ngan, Jenna Gallie, Deepa Agashe

## Abstract

The rate and accuracy of translation hinges upon multiple components – including transfer RNA (tRNA) pools, tRNA modifying enzymes, and rRNA molecules – many of which are redundant in terms of gene copy number or function. It has been hypothesized that the redundancy evolves under selection, driven by its impacts on growth rate. However, we lack empirical measurements of the fitness costs and benefits of redundancy, and we have poor understanding of how this redundancy is organized across components. We manipulated redundancy in multiple translation components of *Escherichia coli* by deleting 28 tRNA genes, 3 tRNA modifying systems, and 4 rRNA operons in various combinations. We find that redundancy in tRNA pools is beneficial when nutrients are plentiful, and costly under nutrient limitation. This nutrient-dependent cost of redundant tRNA genes stems from upper limits to translation capacity and growth rate, and therefore varies as a function of the maximum growth rate attainable in a given nutrient niche. The loss of redundancy in rRNA genes and tRNA modifying enzymes had similar nutrient-dependent fitness consequences. Importantly, these effects are also contingent upon interactions across translation components, indicating a layered hierarchy from copy number of tRNA and rRNA genes to their expression and posttranscriptional modification. Overall, our results indicate both positive and negative selection on redundancy in translation components, depending on a species’ evolutionary history with feasts and famines.

## INTRODUCTION

In the early 1960s, the degeneracy of the genetic code was revealed in the context of multiple synonymous codons encoding a given amino acid. A large body of work since then has uncovered an astonishing degree of redundancy in the translation apparatus. The redundancy is often qualitative, whereby some components can functionally compensate for others. This includes the pool of tRNA molecules that read mRNA codons and deliver the appropriate amino acid during translation. tRNAs with different anticodons may read distinct synonymous codons but carry the same amino acid (tRNA isoacceptors), and are thus functionally degenerate. In addition, tRNA modifying enzymes (MEs) post-transcriptionally alter specific “target” tRNAs, allowing them to read codons that are otherwise decoded by “non-target” tRNAs (Grosjean, 2009). Hence, some non-target tRNAs could be redundant because their function can be carried out by target tRNAs after they get modified. For instance, in *Escherichia coli* the codon CCG can be decoded by the non-target tRNA_CGG_ (encoded by the gene *proK*), but also by the target tRNA_CGG_ after the U at position 34 is modified to cmo^5^U by the cmo modification pathway (in principle rendering the gene *proK* redundant). MEs are thus critical for maximizing cellular decoding capacity (Diwan and Agashe, 2018; Grosjean et al., 2010) and maintaining translational capability requires either a very diverse tRNA pool or the presence of MEs that allow for a compact tRNA set (Diwan and Agashe, 2018; Marck and Grosjean, 2002; Rocha, 2004; Wald and Margalit, 2014). In addition, quantitative redundancy is conferred by large gene copy number (GCN), such that multiple genes can perform identical functions. For instance, bacterial genomes often carry many copies of tRNAs with a given anticodon (tRNA isotypes) (Chan and Lowe, 2009). Similarly, cells typically have many copies of rRNA genes (Roller et al., 2016), which govern ribosome availability (Nomura et al., 1980).Thus, the translation machinery is predicted to be functionally redundant at many levels, with qualitative (multiple codons, tRNA isoacceptors, and MEs) as well as quantitative (GCN) redundancy. However, in many cases the predicted functional redundancy is not experimentally verified, and it remains unclear whether it also influences fitness. We asked: what are the fitness costs and benefits of translational redundancy, and under what conditions do they manifest? Could these costs and benefits explain the evolution of a highly redundant translation apparatus?

In bacteria, selection for rapid growth (often facilitated by nutrient availability) is thought to be an important force that shapes the evolution of many translation components. The maximum possible growth rate is determined by translation efficiency, which in turn depends on concentrations of ribosomes and tRNAs (Ehrenberg and Kurland, 1984; Hu et al., 2020; Kurland and Ehrenberg, 1987; Kurland, C.G., Hughes, 1996). Across species, there are striking positive correlations between maximal growth rate and the number of tRNA and rRNA genes (Dethlefsen and Schmidt, 2007; Ikemura, 1985; Rocha, 2004; Roller et al., 2016; Vieira-Silva and Rocha, 2010; Weissman et al., 2021). Thus, we expect that GCN redundancy in key translation components should be especially beneficial during rapid growth in a nutrient-rich niche. In contrast, under nutrient limitation, expressing redundant genes should be costly because translational output remains constrained by nutrients. However, this overarching growth rate-dependent selection may shape the redundancy of translational components differentially. For instance, selection should have the maximum impact on rRNA, whose GCN shows the strongest correlation with growth rate (Rocha, 2004; Roller et al., 2016) and whose concentrations are most limiting for translation because rRNAs constitute up to 85% of all RNA in rapidly growing *E. coli* (Bremer and Dennis, 1996). During fast growth, cellular rRNA abundance increases by ~250% whereas total tRNA increases only by ~80% (Dong et al., 1996). Consequently, the cellular ratio of tRNAs to rRNAs decreases under rapid growth, indicating greater investment in rRNA (Dittmar et al., 2004). Indeed, deleting rRNA operons in *E. coli* reduces fitness in rich media, but improves fitness in poor media (Condon et al., 1995; Gyorfy et al., 2015; Stevenson and Schmidt, 2004).

In contrast, the growth rate-dependent impacts of redundancy in tRNA GCN remain largely unexplored, with the exception of initiator tRNA genes in *E. coli.* As predicted, in this case the loss of some gene copies is deleterious in rich media and advantageous in poor media (Samhita et al., 2014). However, the growth impact of elongator tRNA GCN is not known. The paucity of data for tRNA redundancy is glaring because bacteria show enormous variation in tRNA pools, driven by evolutionary changes in tRNA GCN as well as MEs (Ayan et al., 2020; Diwan and Agashe, 2018; Saks et al., 1998; Wald and Margalit, 2014). Furthermore, the impacts of redundancy are predicted to vary substantially across different tRNA genes. For instance, the expression of “major” tRNAs that connect frequently used amino acids and codons is more strongly correlated with growth rate (Berg and Kurland, 1997; Dong et al., 1996) and rRNA GCN (Mahajan and Agashe, 2018). The loss of redundancy of these tRNAs should impose a larger fitness cost, a hypothesis that remains untested. Thus, despite strong comparative evidence for growth rate-driven selection, our understanding of the evolution and impacts of redundancy in translation components is far from complete.

While the fitness costs and benefits of redundancy should ultimately be shaped by nutrient availability, redundancy may itself arise via different mechanisms in different translation components, and hence selection may shape the components in distinct ways. For instance, with rRNAs, the loss of GCN redundancy is buffered by strong compensatory upregulation of “backup” gene copies. An *E. coli* strain with deletion of 6 out of 7 rRNA operons can produce about half of the normal levels of rRNA (Asai et al., 1999). As a result, the deletion of a few rRNA genes only moderately reduces growth rate in rich media (Gyorfy et al., 2015), though the direction of the effect is reversed in poor media (see above). Are tRNA pools similarly regulated? While this is possible (e.g. in response to nutrient availability (Fessler et al., 2020; Sørensen et al., 2018), we have no data on compensatory regulation of tRNAs after gene loss. Further, tRNA redundancy is modulated not only by GCN but also by MEs that allow target tRNAs to perform the function of non-target tRNAs. Thus, tRNA gene loss could be buffered by regulation of other tRNA copies, and/or by the action of MEs. Interestingly, fast-growing bacteria tend to have low tRNA diversity (Rocha, 2004), with their decoding capacity likely maintained by the action of multiple tRNA modification pathways (Diwan and Agashe, 2018). Hence, the fitness consequences of tRNA gene loss should be contingent on the availability of ME backups. Conversely, the joint deletion of non-target tRNAs and MEs is predicted to be more costly than the loss of either component alone (Diwan and Agashe, 2018; Wald and Margalit, 2014). Thus, the mechanisms that mediate redundancy as well as the interactions between translation components are important to fully understand the evolution of translational redundancy.

The patterns noted above suggest a hierarchical organization, whereby redundancy in some components and genes is more important than others (e.g. rRNAs vs. tRNAs, and major vs. minor tRNAs). However, as discussed above we have very limited empirical evidence for the fitness consequences of redundancy in different translation components, particularly in the case of tRNA pools. We addressed these gaps by analyzing the nutrient dependent impact of changing redundancy in multiple translational components, alone as well as in combination. Specifically, we tested the following predictions: (1) redundancy in tRNA and rRNA GCN, on the whole, should be important to maintain rapid growth, and the benefits of increased redundancy should be proportional to the achievable growth rate (2) broadly, a reduction in rRNA GCN should have stronger fitness impacts than tRNA GCN (3) across tRNAs, the loss of redundancy should be most impactful for major tRNAs, and for non-target tRNAs when combined with the loss of a relevant ME (4) the fitness impact of reduced redundancy should increase with the severity of the loss, e.g. due to the deletion of multiple gene copies or multiple translational components. We worked with *E. coli* because it has a highly redundant translation machinery (Diwan and Agashe, 2018; Wald and Margalit, 2014) that allowed us to test the impacts of successive losses of redundancy at the level of rRNA genes, tRNA pools, and tRNA modifying enzymes. We first show that, as expected, many components of the translation machinery are indeed redundant with respect to fitness. We then test our predictions by measuring the context-dependent costs and benefits of this redundancy. Our results reveal layered factors that may have shaped the evolution of the translation machinery in bacteria.

## METHODS

### Generating strains

We made all gene deletions in *E. coli* MG1655, which we refer to as the wild type (WT). tRNA deletions were made using Red recombinase, slightly modifying the Datsenko-Wanner method (Datsenko and Wanner, 2000) with longer homology regions of 60–100 bases to increase the probability of recombination. In all but one case (ΔglyVXY) we removed the Kanamycin marker inserted during recombination. We confirmed all strains had marker-less deletions by PCR followed by Sanger sequencing (primers given in Table S1) and Next Generation Sequencing (Illumina HiSeq PE150, >30x coverage). We used P1 transduction to transfer modifying enzyme (ME) deletions received from CGSC (Keio collection) to our WT strain, conducting additional rounds of transduction to make further tRNA deletions as required. Similarly, we combined rRNA deletion strains (from CGSC) with tRNA deletions using P1 transduction. We stored glycerol stocks of each strain at −80°C. Further details are given in the supplementary methods and Table S2.

### Measuring growth parameters

We inoculated strains in LB (Lysogeny Broth, Difco) from individual colonies grown from freezer stocks, and incubated cultures at 37°C with shaking at 180 rpm for 14-16 hours (preculture). For growth rate measurement, we sub-cultured 1% v/v in 48 well microplates (Corning) in the appropriate growth medium: LB, TB (Terrific Broth, Sigma) or M9 minimal medium (M9 salts, 1mM CaCl2, 2.5 mM MgSO4) supplemented with specific carbon and nitrogen sources (“GA”: glucose and cas amino acids, either 1.6% w/v or 0.8% w/v each as specified in the results and figures; or carbon sources alone: lactose 0.05% w/v, pyruvate 0.3% w/v, succinate 0.3% w/v, or glycerol 0.6% w/v). We measured growth rate (*r*) as the change in optical density (OD) read at 600 nm every 20 min or 45 min (for rapid and slow growth respectively), using an automated system (LiconiX incubator, robotic arm and Tecan plate reader). We estimated *r* by fitting exponential equations to OD vs. time curves, using Curvefitter software (Delaney et al., 2013). After 8–12 hours of growth, we estimated the carrying capacity (K) by measuring the maximum OD of late log phase cultures (after 10x dilution in rich media, to accurately estimate ODs higher than 1). We estimated the length of lag phase (L) as the time taken to reach early log phase of growth. This was limited by low temporal resolution, and we were unable to capture differences in L that were smaller than 20 minutes (often observed in the WT and single gene deletions during rapid growth). We estimated relative fitness of each mutant as the ratio of its r, K or L value vs. that of the WT measured in the same experiment.

To measure growth rate under nutrient shifts, we initiated precultures and sub-cultured them as above in a rich medium (TB). From late log phase culture in the rich medium (after 6 hours of growth), we again sub-cultured as above into poor medium (either M9 glycerol or M9 galactose, representing a nutrient downshift). When these cultures reached late log phase in the poor medium (12-16 hrs), we again sub-cultured them back into the rich medium (TB, representing a nutrient upshift). After each transfer (downshift or upshift), we measured growth rate as described above.

### Measuring tRNA pools using YAMAT-Seq

For a subset of our tRNA deletion strains and WT, we measured the relative abundance of tRNAs as described previously (Ayan et al., 2020; Shigematsu et al., 2017). Briefly, we grew three independent replicate cultures of each strain in two media and isolated total RNA from 4 ml (rich medium, LB) or 12 ml (poor medium, M9+0.05% galactose) aliquots of mid-log phase cultures. Next, we carried out a deacylation step to strip amino acids from tRNAs and expose the 3’ deacylated ends. We ligated Y-shaped DNA/RNA hybrid adapters to these ends, and reverse-transcribed ligated products to cDNA. After 11 cycles of PCR-amplification with a proof-reading DNA polymerase, we added sample-specific barcodes, quantified the DNA in each sample, and combined equimolar amounts of all samples. To isolate cDNA corresponding to adapter-ligated tRNAs, we ran the mixture on a 5% native polyacrylamide gel and extracted bands of ~200–280 bp, and extracted DNA from the gel. The purified product was sequenced by the sequencing facility at the Max Planck Institute for Evolutionary Biology (Plön, Germany) using an Illumina NextSeq 550 Output v2.5 kit (Single-end, 150 bp reads). Further details are provided in the supplementary materials.

We sorted raw reads for each sample using exact matches to each unique, 6-bp long Illumina barcode, obtaining a minimum of 707,429 reads per sample of which >99.99% were the expected length (80-151 bp) (Table S3). We assembled each set of reads to the 49 unique reference tRNA sequences predicted by GtRNAdb 2.0 (Chan and Lowe, 2009) for *E. coli* MG1655 (Table S4), allowing up to 10% mismatches, gaps of < 3 bp, and up to five ambiguities per read. We discarded reads that aligned equally well to more than one tRNA sequence. Finally, we *de novo* aligned the unused reads for each sample, and checked the resulting contigs to ensure that none contained substantial numbers of tRNA reads. We calculated the within-sample proportion of reads aligned to each tRNA type and mean mature tRNA isotype proportions for each strain across the three replicates. Finally, we used DESeq2 (Love et al., 2014) in R (version 3.6.0, (Core Team, 2021)) to detect tRNA expression differences between pairs of strains, correcting for multiple testing with the Benjamini-Hochberg procedure (Anders and Huber, 2010). The raw YAMAT-Seq reads and analysis files are available at the NCBI Gene Expression Omnibus (GEO accession number GSE198606) (Edgar et al., 2002).

### Measuring translation elongation rate

We measured translation elongation rate for a subset of our strains, using the native β-galactosidase protein as a reporter as described earlier (Miller, 1972), with some modifications. Briefly, we induced *lacZ* gene expression in actively growing cultures (OD_600_ = 0.5, n = 2–3) with 0.5 mM isopropyl-β-D-thiogalactoside (IPTG). Every 15 seconds, we pipetted out 500 μl culture and immediately mixed it with 100 μl of chloramphenicol (3 mg/ml) to block translation. After 10 mins of incubation on ice, we added 350 μl of Z buffer (reaction buffer) and continued incubation on ice for 1 hour. Next, we added 200 μl of 12 mg/ml ONPG (onitro-phenyl galactopyranoside, substrate for β-galactosidase). After 1–1.5 hour of incubation at 30°C to allow the full development of colored product (o-nitrophenol) due to enzyme activity, we stopped the reaction by adding 500 μl of 1M Na_2_CO_3_. After a brief centrifugation step to collect debris (5000 g, 1 min), we transferred the supernatant to a 96 well microplate to assay the formation of o-nitrophenol by measuring OD_420_. We converted OD values to Miller Units (MU) as per the original protocol, and from a plot of Miller Units (MU) of β-galactosidase vs. time, we estimated the first time point showing an increase in MU (after induction) as the time taken to synthesise one molecule of β-galactosidase. The elongation rate (in amino acids per second) was inferred by dividing the length of the β-galactosidase protein (1019 amino acids) by this time.

## RESULTS

### Altering redundancy in translation components

Prior work demonstrates the functional redundancy of some bacterial translation components with respect to translation rate or accuracy. However, it remains unknown whether and under what conditions this functional redundancy translates into fitness consequences. The genome of *E. coli* MG1655 (wild type, WT) encodes 42 tRNA isotypes with varying copy number (total 86 tRNA genes) (Chan and Lowe, 2016), 5 tRNA modification pathways (Diwan and Agashe, 2018) that modify the 34th base of the tRNA or first base of the anticodon, and 7 rRNA genes in distinct operons (Quan et al., 2015). We reduced redundancy in translation components in three ways (Fig 1, Table S2). (1) We generated 23 distinct mutant strains of WT that represented a total of 28 deleted tRNA genes, with 20 strains carrying single tRNA deletions and 3 strains carrying multiple tRNA deletions. These strains denoted a direct genomic loss of redundancy, potentially altering the cellular tRNA pool (sets I, II and III, Fig 1, Table S2). (2) Post-transcriptional modification enhances wobble pairing by adding anticodon loop modifications to “target” tRNAs. Hence, non-target tRNAs are made redundant by modified tRNAs. To reduce this form of redundancy, we deleted key enzymes (MEs) within four tRNA modification pathways of WT (set IV, Fig 1, Table S2), as well as some of their non-target tRNAs (set V, Fig 1, Table S2) and a target tRNA in one case. (3) Finally, to lower redundancy in rRNA genes, we used strains carrying 1–4 rRNA operon deletions, including deletions of interspersed tRNA genes (Quan et al., 2015) (set VI, Fig 1, Table S2). In one of the strains missing four rRNA operons, we made additional tRNA deletions, so that both tRNA and rRNA would be limiting (set VII, Fig 1, Table S2). Overall, we used 43 mutant strains covering 15 amino acids, 33 tRNA genes, 3 tRNA modifying systems, and 4 rRNA operons (Fig 1, Table S2).

**Figure 1:**
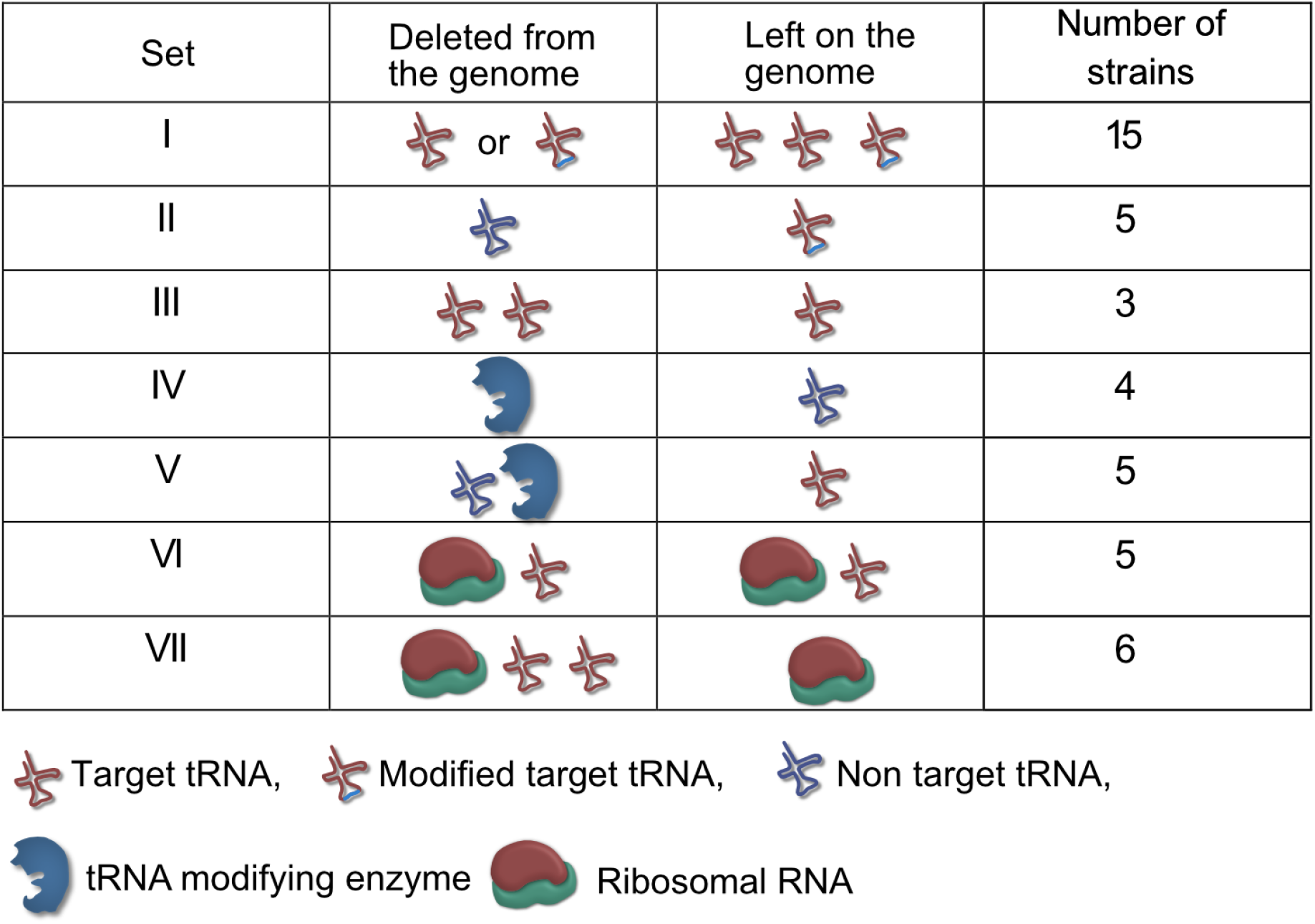
Summary of experimental manipulation of redundancy in translational components. Target tRNAs indicate tRNA that are post-transcriptionally modified by a tRNA modifying enzyme, allowing them to perform the function of specific non-target tRNAs. The symbols represent qualitative differences across sets rather than the exact number or diversity of redundant components. Further details of strains in each set are given in Table S2.

### The loss of tRNA redundancy has highly variable growth impacts

Of our strains, 15 represented the deletion of single tRNA genes. All but one *(proL)* were predicted to be redundant because they have other gene copies encoding the same tRNA isotype (set I in Fig 1, Table S2). In some cases the original GCN was small, so that our manipulation left a single redundant copy *(phe U/V* and *ser W/X).* In five other strains we deleted single tRNA genes that appear redundant because MEs should allow other (target) tRNAs to perform their function (set II, Table S2). Given the expected functional backups for deletions in sets I and II, we predicted that the loss of redundancy should have relatively weak fitness consequences. Indeed, in complex rich media (LB and TB), the growth rate of these strains was largely similar to the WT, with the highest impact representing ~15% change in growth rate (R_rel_ values between 0.85 and 1.15; R_rel_ is the ratio of mutant growth rate to WT growth rate, so R_rel_ > 1 indicates faster growth of mutant). Only 10 of 20 strains showed a significant difference in at least one of the complex rich media, 6 with faster growth and 4 with slower growth than WT (sets I and II in Fig 2A, Fig S1A, Fig. S2A, Table S5). Interestingly, deleting the only genomic copy of *proL* had negligible effects on growth (Fig 2A), possibly because the relevant codon is used very rarely (Table S2). On the other hand, deleting different gene copies encoding the same tRNA isotype (e.g. *asnT* vs. *asnV,* each encoding tRNA-Asn(GUU)) impacted growth differently, corroborating previous reports of functional differences between tRNA copies (Dittmar et al., 2004). As predicted, a more severe reduction in redundancy via deletion of multiple tRNA gene copies (leaving only one backup copy of many, set III, Fig 1) reduced growth rate substantially, with a mean R_rel_ of ~0.75 (i.e. ~25% change) (Fig 2A, Fig S1A, Fig S2A). Over half of the 20 strains also showed a significant difference in the length of the lag phase, though the direction of the effect varied across strains and media (Fig S2B, Fig S3A). Consistent with the growth rate results, set III strains showed the maximum increase in lag phase length. However, barring a few exceptions, the loss of tRNA redundancy had negligible impact on growth yield regardless of the severity of the manipulation (Fig S2C, Fig S4A).

**Figure 2:**
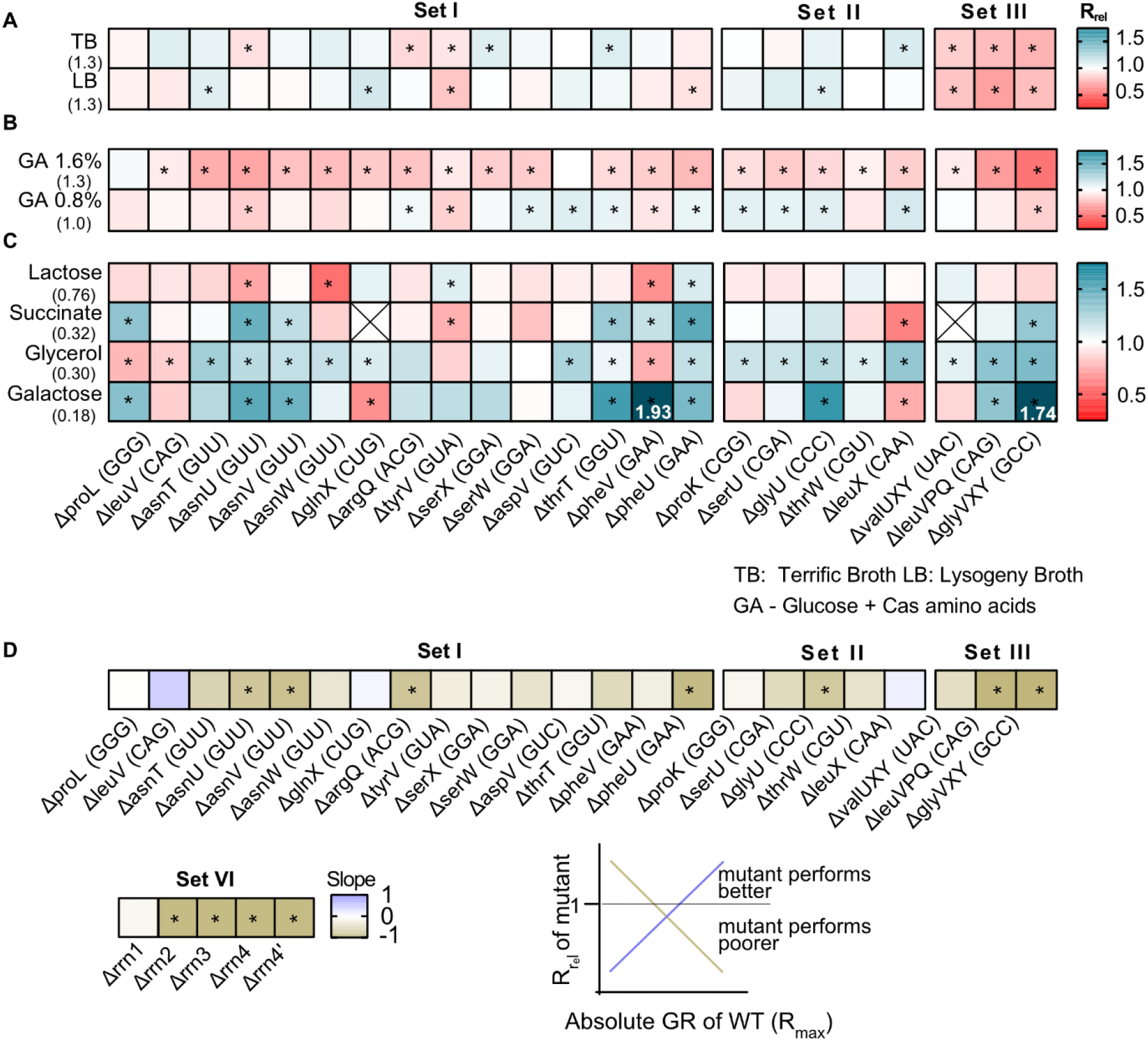
The fitness impact of loss of tRNA redundancy varies with nutrient availability. (A–C) Heat maps show the relative growth rate of tRNA gene deletion strains (“mutant”) (R_rel_=GR_mutant_/GR_WT_) in different growth media (see Table S5 for statistics). The anticodon of each deleted tRNA gene is indicated in parentheses on the x-axis. The absolute exponential growth rate (doublings/hour) of WT in each medium is indicated in parentheses on the y-axis. Box colors indicate the impact of each gene deletion (red: R_rel_ <1, mutant grows more slowly than WT; blue: R_rel_ >1, mutant grows faster than WT; n = 3–4 per strain per medium); statistically significant differences from WT are indicated by asterisks (ANOVA with Dunnet’s correction for multiple comparisons). Boxes with an “X” indicate cases where strains failed to grow exponentially. Panels show R_rel_ in (A) complex rich media (B) permissive rich media (M9 salts supplemented with indicated concentrations of glucose and cas amino acids) (C) poor minimal media (M9 salts) with the indicated carbon source but no cas amino acids. (D) For each mutant we estimated the Spearman’s rank correlation between the growth rate impact of the gene deletion (R_rel_) and the respective maximal WT growth rate (R_max_) across 8 growth media (Fig S5). The heat map shows the slope of this correlation for each mutant. Darker colors indicate a stronger negative relationship, i.e. a higher cost of redundancy in poor media. Statistically significant non-zero slope values are indicated by asterisks. Data for mutants with rRNA operon deletions (across 6 growth media) are also included in this panel (see Fig 6B).

Next, we tested the fitness of tRNA deletion strains in more permissive rich media with easy-to-use sources of carbon (glucose) and amino acids (casamino acids) (“GA”), where the WT growth rate is similar to that in complex rich media. Here, tRNA loss was uniformly deleterious, with 18 of 20 strains showing significantly slower growth (GA1.6, sets I and II, mean R_rel_ = 0.84 and 0.88 respectively; Fig 2B, Fig S1B, Fig S2A, Table S5). Reducing the glucose and casamino acid concentration reversed this effect (GA0.8, sets I and II, mean R_rel_ = 0.98 and 1.02 respectively; Fig 2B), suggesting that nutrient availability determined both the direction and uniformity of the impact of tRNA redundancy. As with complex rich media, strains with a severe loss of redundancy tended to show the largest reduction in fitness (set III, Fig 2B, Fig S1B, Fig S2A). However, the impacts on other growth parameters were more variable. In GA1.6, 6 of 20 strains (sets I and II) showed a significantly shorter lag phase than WT while 5 showed a longer lag phase; but in GA0.8, only 1 strain had a longer lag phase and 12 strains entered the exponential growth phase faster than WT (Fig S3B). Concomitantly, we observed little change in the growth yield of these strains, with only 3–4 strains showing a significant difference in either medium (Fig S4B). Unlike the patterns in complex rich media, set III strains did not show stronger effects on either lag phase length or growth yield (Fig S2B–C). Overall, in media where easily accessible nutrients are plentiful, even a small loss of tRNA redundancy strongly hindered rapid growth but had weak and/or inconsistent effects on the lag phase and growth yield.

Finally, we measured fitness in poor media where nutrients should severely limit translation, and maintaining tRNA redundancy may be costly. When using lactose (which reduces WT growth rate to ~50% of LB), the loss of redundancy had a weak and variable impact, with only 5 of 23 strains (across sets I–III) showing a significant difference from WT (Fig 2C, Table S5). However, in poorer media containing succinate, glycerol or galactose (where WT growth rate is reduced to ~10–25% of LB), tRNA deletions were often beneficial (7, 15 and 9 out of 23 strains respectively) and only 2–3 strains had slower growth than the WT (sets I–III, Fig 2C, Fig S2A). Again, set III strains tended to show the maximum benefit of tRNA loss (Fig 2C, Fig S2A). Although the impacts on other growth parameters varied across strains and growth media, most strains had a shorter lag phase (20 of 23 strains) and a higher yield (14 of 23 strains) in at least one poor medium (Fig S2B–C, Fig S3C, Fig S4C). Hence, the loss of redundancy appeared to be generally beneficial in poor media.

Overall, these results indicated that a severe loss of tRNA redundancy amplified the fitness impacts of tRNA deletion, but the magnitude and direction of the effects varied substantially across growth media. However, in all growth media, the fitness impacts were generally similar for sets I and II (Fig S2A–C), indicating that the nature of the backup available to maintain tRNA pools (redundant gene copies vs. ME activity) does not alter the impact of tRNA loss.

### tRNA redundancy is beneficial during rapid growth but costly under nutrient limitation

The results above showed that the fitness impacts of tRNA loss depend qualitatively on the growth medium. To test whether these patterns are quantitatively explained by growth limits set by nutrient availability, for each engineered strain we estimated the relationship between the relative impact of loss of redundancy (R_rel_) and the maximum attainable WT growth rate (R_max_), across all growth media tested (data from Figs 2A–C). Since WT has the highest level of redundancy and presumably the weakest internal limits on translation rate, we expected that the WT R_max_ reflects the nutrient capacity of each growth medium (i.e. externally placed limits on growth). Of 28 strains (including 5 rRNA operon deletions described later in the results section), 26 showed a negative correlation between R_max_ and R_rel_, with a significant relationship in 11 cases (Fig 2D, Fig S5). Thus, tRNA and rRNA loss tends to be more beneficial (i.e. redundancy is more costly) under conditions of low nutrient availability, when the maximum possible growth rate is constrained.

This pattern was further supported by experiments performed during nutrient shifts, where cultures in exponential growth phase were transferred from rich to poor media and vice versa. Note that this setup differs from the previous growth measurement (Fig 2) where late stationary phase cultures grown overnight were transferred to either rich or poor media. Of the 27 tRNA and rRNA deletion strains tested, all but one had a growth rate that was comparable to WT (16 strains) or higher than WT (10 strains) after transitioning from rich to poor media (i.e. during a nutrient downshift, note data distribution along the x-axis in Fig 3; Table S6). In contrast, after a nutrient upshift, 11 strains showed significantly slower growth in one or both pairs of media, and only 2 showed a significantly faster growth than WT (note data distribution along the y-axis in Fig 3; Table S6). Thus, gene loss is beneficial during a nutrient downshift but deleterious in a nutrient upshift. These patterns were also most consistent for strains in sets III and VI (described later in the results section), corroborating our results from constant environments where we observed large impacts of redundancy in the same strains. Strains in the bottom right quadrant of Fig 3 are especially interesting because they represent cases where the loss of redundancy is beneficial in a nutrient downshift but deleterious in a nutrient upshift. Hence, these genes should be important when ramping up translation in a nutrient-rich environment. In this category, we observed one strain each from sets I and II, and 4 strains from set VI (Table S6). Thus, redundancy in tRNA genes can be beneficial during rapid growth, but is generally costly in poor media where nutrients are limited.

**Figure 3:**
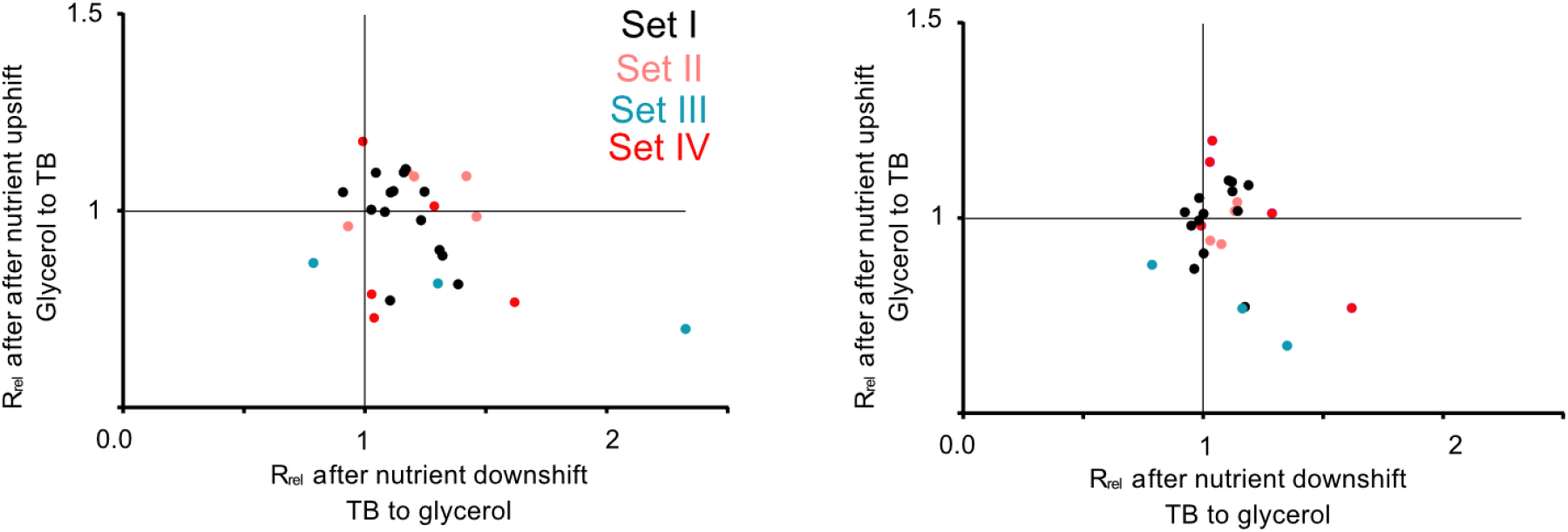
Redundant tRNA copies are beneficial during nutrient fluctuations. Relative growth rate of mutants (sets I to IV, Fig 1) upon a nutrient downshift (R_rel_ on the x-axis) and an upshift (R_rel_ on the y-axis). R_rel_ was calculated relative to WT (growing in identical conditions), as described in Fig 2. Strains were transferred between a rich medium (TB, terrific broth) and a poor medium (M9 salts + 0.6%glycerol or 0.05% galactose). Each data point represents the mean R_rel_ for a mutant (n = 4), with the color indicating the set to which it belongs (Fig 1). Mutants in the bottom right quadrant perform better under a nutrient downshift (i.e. when moving from a rich to poor medium), but more poorly than WT under a nutrient upshift (i.e. moving from a poor to rich medium). Mutants in the top right quadrant perform better than WT in both types of nutrient fluctuations. Statistics are reported in Table S6.

### Gene regulation cannot compensate for loss of tRNA gene copies

Recall that when tRNA redundancy was lowered to an extreme (set III, Fig 1), cells still had at least one copy of each gene and could potentially compensate for tRNA loss by upregulating this backup gene. However, these strains paid a substantial fitness cost in rich media (Figs 2A–B, Fig S1A, Fig S2A), suggesting that such upregulation could not fully compensate for severe tRNA gene loss. Conversely, the large fitness benefit of losing the same tRNAs in poor media (Fig 2C) suggests that these genes are not sufficiently downregulated in poor media, with cells paying a maintenance cost. To test these hypotheses, we measured tRNA expression levels in WT and four tRNA deletion strains in a rich (TB) and a poor medium (M9 galactose), focusing on strains from sets II and III where we observed strong fitness effects. In the WT, 26 of 42 tRNAs did not show a significant difference in expression across media, confirming minimal regulation (Fig 4A, Fig S6). In fact, 10 tRNAs were significantly upregulated in the poor medium relative to the rich medium (top row, Fig 4A; Table S7). In contrast (and as expected), all tested tRNA deletion strains had lower expression of focal tRNA isotypes in the rich medium (Fig 4B, left panel), showing that the backup gene copies are not upregulated sufficiently to compensate for the loss of deleted tRNAs. Even in the poor medium, WT continued to express more of the focal tRNAs compared to the respective deletion strains (right panel, Fig 4B; Fig S7). Hence, compared to gene regulation, change in gene copy number allows a stronger (in this case, also more beneficial) response to the nutritional environment.

**Figure 4:**
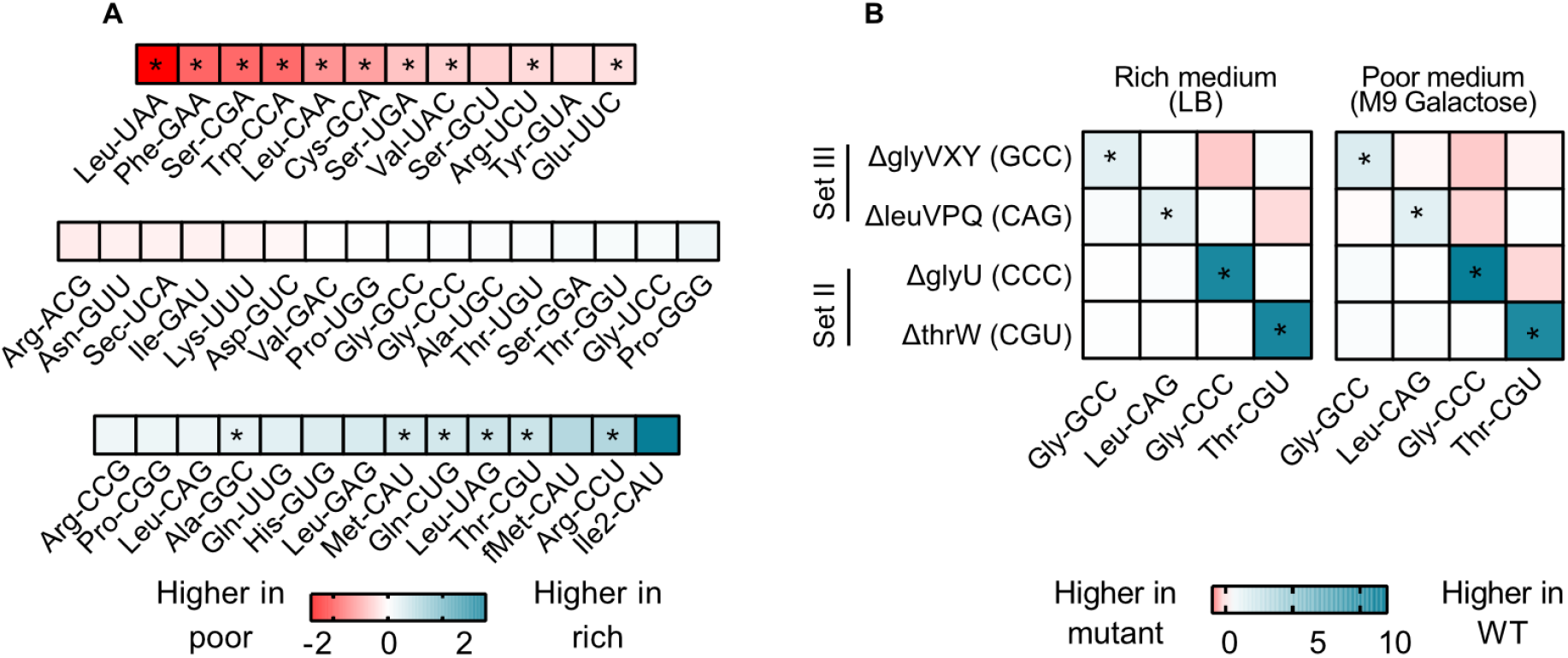
Gene copy number changes confer stronger control over tRNA expression than gene regulation. tRNA expression levels were measured for WT and 4 tRNA deletion mutants in a poor medium (M9 + 0.05% galactose) and a rich medium (LB) (n = 3 per medium per strain) using YAMAT-Seq. Relative expression across strains or across media was calculated as mean log2 fold change. Significant differences are shown with asterisks (pairwise Wald tests with Benjamini-Hochberg correction for multiple comparisons; Table S7). (A) Expression level of all 42 WT tRNA isotypes in poor medium relative to rich medium. Isotype is indicated on the x-axis. Blue indicates higher expression in rich medium and red indicates higher expression in poor medium. (B) The impact of tRNA gene deletion on the expression of focal isotype tRNAs (indicated on the x-axis), in WT vs. mutant strains (tRNA gene deletions indicated on the y-axis), in a rich and a poor medium. Darker blue colors indicate lower expression in mutant, and darker red indicates higher expression compared to WT.

Interestingly, in the tRNA deletion strains the expression of many other tRNA species differed from the WT expression level. YAMAT-Seq measures relative tRNA levels, so that when we deleted highly expressed tRNAs (e.g. ΔglyVXY and ΔleuVPQ) we would automatically alter the relative expression levels of many other tRNAs. However, the difference between WT and tRNA deletion strains varied systematically across media (Fig S7) and was also evident with the deletion of intermediately expressed tRNAs. Hence, these differences likely reflected transcriptional regulation or processing. In other words, the transcriptional response to the loss of tRNA gene copies was stronger in rich medium, even though this regulation did not restore fitness completely. This further indicated that in rich media, gene copy number is limiting, whereas in poor media nutrients are a major limiting factor. Note that while the deletion of non-target tRNAs (i.e. ΔthrW and ΔglyU, with ME backup) resulted in reduced expression of the deleted tRNA as expected (Fig S7), these gene deletions had only weak fitness effects (Fig 2). Thus, MEs indeed serve as a backup and render non-target tRNAs redundant. Overall, these data confirmed that moderate to severe loss of redundancy at the gene copy level is not fully rescued by regulation of backup gene copies.

### Redundant tRNAs do not contribute to translation when nutrients are limited

We showed above that the WT over-produces many tRNAs in poor media, potentially explaining its low fitness relative to the tRNA deletion strains. We suspected that although in rich media such “surplus” tRNAs contribute to translation, this may not happen in poor media where growth is limited by nutrient availability rather than translation efficiency, and levels of charged amino acids drop (Dittmar et al., 2005; Elf et al., 2003). Thus, in rich media, the loss of tRNAs should decrease translation; but in poor media, this effect should be weak. We therefore estimated translational output in a subset of our strains, by measuring the translation elongation rate of the native beta-galactosidase protein during the log phase of growth. As predicted, in a rich medium (LB) all strains with low redundancy had a significantly slower elongation rate than WT (Fig 5). In a permissive medium (GA), elongation rates were usually not significantly different from WT, a pattern that was also observed for another reporter protein (GFP, Fig S8). However, in a poor medium (M9 glycerol), elongation rates were often higher than WT (Fig 5), indicating that the loss of tRNA genes had a net beneficial impact on translation elongation. Again, the effect of tRNA deletion on translation rate increased with the magnitude of the loss of redundancy, with set III strains showing the largest effect size (Fig 5, Fig S8). Overall – as expected from the correlation between growth rate and translation rate – these results mirror the impacts of tRNA redundancy on fitness (Fig 2). Thus, under nutrient limitation, redundant tRNAs are expressed (with cells paying the cost of expression), but these tRNAs do not contribute to growth because they do not increase translational output enough to compensate for the cost of expression.

**Figure 5:**
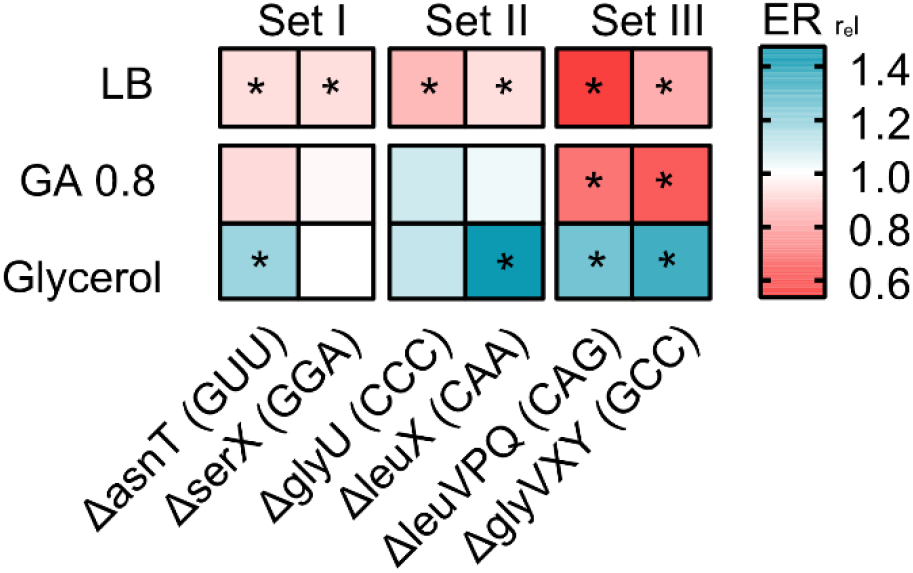
Loss of redundant tRNAs decreases translation output during rapid growth, but increases translational output in poor media. The heat map shows the translation capacity of tRNA deletion mutants in rich (LB), permissive (M9 + 0.8% GA) and poor medium (M9 + 0.6% glycerol), measured as the relative protein elongation rate (ER, increase in the length of ß-galactosidase protein per unit time after induction; ER_rel_= ER_mutant_/ER_WT_; n = 2 per strain per medium). Red indicates lower reporter protein production in the mutant per unit cell density (i.e. reduced translation capacity), and blue indicates increased translation capacity relative to WT. Significant differences are indicated with asterisks (ANOVA with Dunnet’s correction for multiple comparisons).

### Loss of redundancy in multiple translation components reveals layered fitness impacts

Recall that tRNA deletion strains with severely reduced redundancy (set III, with only 1 gene copy remaining) showed stronger fitness effects than strains in set I and II. Further, strains in set II had similar fitness as those in set I, potentially because the fitness impact of the tRNA deletion (which also reduced the levels of the deleted tRNA isotype) was masked by the presence of MEs. Thus, the fitness impact of redundancy generally increases with the magnitude of the loss, as predicted by comparative evidence across genomes; but this has not been explicitly demonstrated. To do so, we further lowered translational redundancy by simultaneously deleting multiple translation components.

Loss of the modifying enzyme-coding genes *cmoA, cmoB, mnmG,* and *tgt* (set IV, Fig 1) significantly reduced growth rate in all media (Fig 6A, Table S5). Consistent with our prediction, the combined fitness effect of ME deletion and non-target tRNA deletions (set V, Fig 1) was stronger than the effect of deleting only the non-target tRNA genes, with all 5 tested co-deletions showing a significantly higher effect in at least one medium (Fig 6A). However, the impact of co-deletions was statistically indistinguishable from the effect of deleting MEs alone, in all except the leucine triple deletion in LB and Glycerol (Table S5). Conversely, and as expected, co-deletion of the ME *tgt* and its target tRNA (asnU) had a significantly different impact from the deletion of the target tRNA alone in all media. This suggested that the effect of co-deletion is largely driven by ME loss, provided that the co-deleted tRNA is not a target tRNA. As observed with tRNA gene deletions (Fig 2), in rich media the loss of ME redundancy tended to be deleterious, whereas in poor media the effects were more variable and included cases where gene loss was beneficial. Note that the joint importance of redundancy in ME and non-target tRNAs is observed in both rich and poor media, with more instances of beneficial effects in the latter. Thus, these results confirmed that MEs serve as important backups when the diversity of the tRNA pool is depleted and that a reduction in redundancy (via tRNA GCN and/or ME loss) is generally beneficial in poor media.

**Figure 6:**
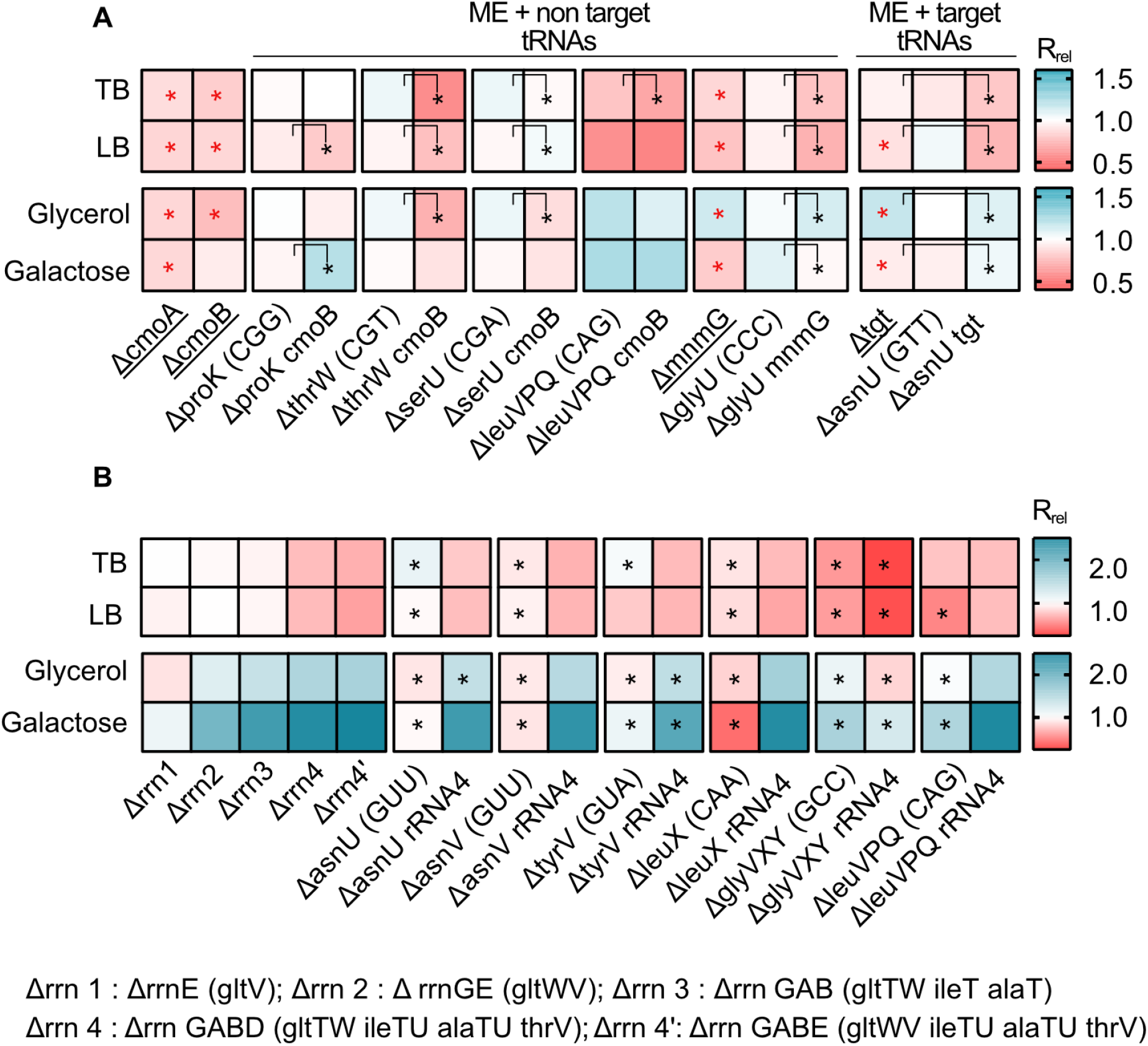
Impacts of manipulating redundancy in multiple translation components are highly variable. Impact of tRNA, rRNA and modifying enzyme (ME) gene deletion on growth rate in different media, as described in Fig 2. Box colours indicate the impact of gene deletion on growth rate relative to WT (red: R_rel_ <1, mutant grows more slowly than WT; blue: R_rel_ > 1, mutant grows faster than WT; n = 3–4 per strain per medium). Panels show R_rel_ for (A) co-deletion of MEs and tRNAs. The anticodon of each deleted tRNA gene is indicated in parentheses on the x-axis. MEs were either deleted alone (underlined strains) or co-deleted with the respective non-target and target tRNAs. ME deletions strains were compared with the WT to establish statistically significant differences, indicated with a red asterisk. For co-deletions of MEs and non-target tRNAs, comparisons are shown between non-target tRNA deletion and ME+non target tRNA codeletion (as indicated by the asterisks and vertical square brackets). For codeletions of MEs and target tRNA, comparisons are shown between ME deletions and ME+ target tRNA codeletion (as indicated by the asterisks and vertical square brackets). Other statistical comparisons, not shown in the figures are in Table S5. (B) co-deletion of rRNAs and tRNAs. rRNA operons are abbreviated as shown in the key; tRNA genes that were deleted as part of the operon are indicated in parentheses. Additional tRNAs (outside the rRNA operon) that were deleted in combination with rrn4 are indicated on the x-axis. Asterisks indicate statistically significant impacts (ΔrRNA4 vs. ΔrRNA4+tRNA deletion, (ANOVA with Dunnet’s correction for multiple comparisons), Table S5).

Next, we tested the combined effect of altering redundancy in rRNA and tRNA genes. As noted earlier, deleting rRNA operons simultaneously removes some tRNA genes located in the operon, so that even a single rRNA operon deletion is effectively a co-deletion. However, all the tRNAs deleted in this manner had multiple genomic backup copies (Table S2). Given the strong regulatory compensation of rRNA loss observed in prior work (Asai et al., 1999; Elf et al., 2003; Quan et al., 2015), it was not surprising that the deletion of up to three rRNA operons (along with up to 4 tRNA genes) had a very weak impact on growth rate in rich media. However, in poor media the fitness impact was evident even with the loss of only two rRNA operons (and 3 tRNA genes). Mimicking the patterns observed for specific tRNA deletions, a more severe loss of rRNA and tRNA redundancy (from 1 to 4 rRNA operon deletions, set VI, Fig 1, Table S5) was detrimental in rich media (set VI, Fig 6B) and increasingly beneficial in poor media (set VI, Fig 6B).

Further loss of redundancy – via simultaneous deletion of 4 rRNA operons (including 7 tRNA genes) and extra-operonic tRNA genes (set VII, Fig 1, Table S2) – had mixed effects on growth rate. While in every case, the impact of the rrna4 deletion was significantly greater than the tRNA deletion alone (Table S5), the effect of tRNA co-deletions varied across media. In rich media, only 1 of the 6 cases tested showed a significant additional fitness reduction upon co-deleting rRNA and extra-operonic tRNA. However, in poor media, tRNA loss led to an additional fitness benefit in 3 out of 6 co-deletions. Thus, when growth is strongly limited by the availability of rRNA (and hence mature ribosomes), lack of tRNA is less detrimental to translation and growth rate. Conversely, when growth is limited by both nutrients and rRNAs, shedding tRNAs appear to be additionally beneficial.

Together, our results lead to the following conclusions. First, we find that simultaneous deletion of multiple copies of tRNA genes or rRNA genes has more severe fitness consequences than the loss of single gene copies. Second, increasing the severity of reduction in translational redundancy via co-deletion of MEs and tRNAs amplifies the fitness consequences of losing redundancy. Lastly, when nutrients are unlimited, rRNA becomes limiting and the loss of tRNAs has little additional impact on fitness; but when nutrients are limiting in the first place, shedding different translation components (rRNA and tRNA) additively increase fitness.

## DISCUSSION

The process of protein synthesis is central to life, and is especially important to understand bacterial evolution given the direct link between translation, growth rate and fitness. Translation rate is affected by several genomic (eg. tRNAs, rRNAs, tRNA modifying and charging enzymes) and environmental components (eg. nutrient availability). Comparative analyses show that the genomic components have different degrees of redundancy across taxa, and suggest that this redundancy (in the fitness context) should be shaped by the strength of ecological selection for rapid growth (Rocha, 2004; Roller et al., 2016). While lateral gene transfer and gene deletion/duplication make translation components labile even in the absence of selection, the correlation between maximum growth rate and redundancy of tRNA or rRNA GCN suggests a strong role for selection (Rocha, 2004; Roller et al., 2016), likely imposed by nutrients from the environment. Hence, nutrients ultimately limit translation. Cells can meet this environmental limit and maximize fitness by shaping genomic factors to achieve the maximum attainable translational output. Such modulation of the cellular machinery should occur at physiological (short-term) as well as evolutionary timescales. For instance, cells can control translation rates via rapid regulation of translation components (Wilusz, 2015), including via degradation of ribosomes and tRNAs during nutrient starvation (Fessler et al., 2020; Sørensen et al., 2018). Across-species patterns of rRNA and tRNA GCN are consistent with such selection, as discussed in the Introduction. Similar arguments can also be made for redundancy across other translation components. Together, this suggests that the environment sets the limits of translation, according to which natural selection shapes the genomic redundancy of translational components. However, empirical evidence for a common underlying selection pressure shaping redundancy across various translational components has been missing.

Here, we provide such evidence, showing that several components of the *E. coli* translation apparatus are indeed functionally redundant, and that the costs and benefits of this redundancy vary with nutrient availability. When nutrients permit rapid growth, the loss of redundancy in both tRNAs and rRNAs is detrimental, as these components become limiting for translation. This is especially true for multiple deletions of abundantly used components such as rRNAs, MEs that modify many different tRNAs, and frequently used tRNAs (such as for glycine and leucine). These results support prior predictions of larger fitness consequences following the loss of major tRNAs that read abundant codons and respond strongly to fast growth (Dong et al., 1996), or tRNAs that make larger contributions to the tRNA pool (Bloom-Ackermann et al., 2014; Kanaya et al., 1999). The observed variability in fitness impacts across tRNAs is therefore at least partially explained by the relative use of different codons. Our results also support the broad prediction that high tRNA levels should be most critical during rapid growth (Mahajan and Agashe, 2018; Vieira-Silva and Rocha, 2010), when the correlation between tRNA levels and gene copy number is strongest (Dong et al., 1996). Conversely, when nutrients are limiting, rRNA as well as tRNA gene loss is beneficial. Importantly, we show that the expression cost of high translational redundancy of WT *E. coli* is not met by increased translation rate in poor media, and imposes a substantial fitness cost.

While these observations suggest that nutrient availability can guide the evolutionary optimization of the cost to benefit ratio of the bacterial translation machinery, interactions (and potential hierarchies) amongst different components suggest multiple routes of optimization. We observe that the loss of rRNA genes is generally more impactful than the loss of tRNAs; and when rRNAs are limiting, the additional loss of tRNAs has a relatively weak effect. We suggest that this is because rRNAs set the first internal limit on translation rate, as predicted by prior work (see the Introduction). However, further experiments are necessary to separate the independent impacts of rRNA and tRNA genes linked within operons. Interestingly, even in complex bacterial communities, addition of extra resources enriched for taxa with more rRNA copies (Wu et al., 2017) and the growth rate response to nutrient addition was positively correlated with rRNA copy number (Li et al., 2019). The next internal limiting factor for translation appears to be available tRNA pools, determined by the nested impacts of tRNA gene copy number, transcriptional regulation of gene expression, and MEs. Overall, tRNA gene copy number has a stronger impact on translation and fitness than the regulation of different isotype copies, corroborating prior results with yeast (Percudani et al., 1997). However, in the special case of non-target tRNAs, the relevant MEs have a stronger impact than the tRNA genes. This is not surprising given the predicted functional redundancy between MEs and non-target tRNAs (Diwan and Agashe, 2018; Grosjean et al., 2010). As a result of this layering, a severe loss of tRNA redundancy becomes important for fitness only when many copies of an isotype are lost, or when a non-target tRNA is co-deleted with a relevant modifying enzyme. Conversely, these results predict that an increase in redundancy (e.g. due to occupation of a more nutrient-rich niche and selection for rapid growth) may occur via increasing isotype gene copy numbers, increasing non-target tRNAs, or gaining a relevant modifying enzyme. These predictions from our data from *E. coli* closely match the patterns observed in comparative analyses across bacteria: selection for rapid growth is strongly correlated with gene copy number (Eduardo P.C. Rocha, 2004), and lineages may lose either MEs or non-target tRNAs, but not both (Diwan and Agashe, 2018).

Together with prior work, our results also suggest that fast-growing organisms such as *E. coli* have evolved to rely strongly on gene copy number to maximize translation rates. Presumably, such species are either able to bear the costs of maintenance of surplus tRNA genes during periods of slow growth, or across longer evolutionary timescales the extra tRNA copies provide a net benefit. We therefore suggest a model whereby bacterial growth rate is primarily limited by external nutrient availability, then by rRNA molecules, and finally by tRNA pools (determined by tRNA GCN and MEs, and secondarily via tRNA gene regulation). Thus, the layered costs and benefits of high translational redundancy – both within and across distinct components – are ultimately determined by the environmental context. Importantly, this model predicts that prolonged selection for rapid growth should cause successive evolutionary changes in the redundancy of different translation components, with the general order of events determined (and parallel routes offered) by the hierarchical layering of components.

We hope that future work will test this model and enrich it by considering additional translational components and factors that may drive their evolution. For instance, genome GC content is strongly associated with tRNA GCN and diversity across the bacterial phylogeny (Diwan and Agashe, 2018; Wald and Margalit, 2014), as well as with codon bias (Hershberg and Petrov, 2010). An understanding of the ecological and evolutionary pressures that drive shifts in GC content would thus be useful to understand the impact of selection for rapid growth on genome GC. It is also worth considering other evolutionary processes that can alter redundancy, such as genetic drift that may facilitate the loss of MEs (Diwan and Agashe, 2018). In addition, here we have focused on selection on translation rate, primarily tested using growth rate. However, our results hint at interesting effects of the loss of redundancy on other growth parameters such as yield and lag phase that may be orthogonal to growth rate. These measurements had low resolution in this study, but explicit and better analysis of such impacts may reveal the effects of selection in niches where yield or survival (rather than growth rate) determine fitness. Finally, we note that selection may act via other cellular functions performed by some translational components (Shepherd and Ibba, 2015), or on translational accuracy (Gingold and Pilpel, 2011). While global mistranslation can provide fitness benefits in stressful contexts (Jones et al., 2011; Samhita et al., 2020), the nature and strength of selection acting on translational accuracy and the relationship between translation rate and accuracy remains poorly understood (Drummond and Wilke, 2009). Nonetheless, prior work suggests that both tRNA pools and MEs influence translation accuracy (Manickam et al., 2016) and protein aggregation (Fedyunin et al., 2012). Thus, selection for accuracy could also shape the evolution of tRNAs and MEs. We therefore suggest that expanding the layers of organization of translational components in our model will prove fruitful in gaining a deeper understanding of the evolution of translational redundancy.

In summary, our experiments demonstrate that several components of the translation machinery are redundant in *E. coli*, the costs and benefits of which vary based on nutrient availability – an environmental variable that likely shaped the redundancy in the first place. Our results support the broad idea that translational limits imposed by different components and their interactions generate multiple translational optima and make many paths feasible (Grosjean et al., 2014; Higgs and Ran, 2008), depending on the selective context (Eduardo P.C. Rocha, 2004). We propose a model with hierarchies of translation components that may allow maximization of translational output and fitness. It remains to be seen to what extent these hierarchies operate across diverse taxa, and which of the possible parallel routes are taken during the course of bacterial evolution in nature. Further studies on the layers and hierarchies connecting translational components will shed light on the molecular toolkits underlying evolutionary transitions between slow vs. rapid translation.

## ACKNOWLEDGEMENTS

We thank Joshua Miranda, Laasya Samhita, Shyamsunder Buddh and Umesh Varshney for discussion and comments on the manuscript; Joshua Miranda, Laasya Samhita, Mrudula Sane and the NCBS NGS facility for help with genome sequencing; Gaurav Diwan and Joshua Miranda for setting up and maintaining our automated growth measurement system; Gunda Dechow-Seligmann for helping with data collection for YAMAT-seq; and the NCBS laboratory kitchen staff for their crucial support overall and especially during the COVID-19 pandemic. We acknowledge funding and support from the National Centre for Biological Sciences (NCBS-TIFR) and the Department of Atomic Energy, Government of India (Project Identification No. RTI 4006) to DA, CSIR-UGC-NET June/2018/430 fellowship to PKR, the Max Planck Society (JG and WYN), and the International Max Planck Research School for Evolutionary Biology (WYN).

## AUTHOR CONTRIBUTIONS

PKR conceived and designed the study, conducted experiments, and analysed data. WYN designed and conducted the YAMAT-Seq experiments. JG designed, conducted and analysed the YAMAT-Seq experiments, and acquired funding. DA conceived and designed the study, directed experiments and analyses, and acquired funding. DA and PKR wrote the paper with input from JG and WYN.

