## Supplementary Information for "The layered costs and benefits of translational redundancy"

### SUPPLEMENTARY METHODS

#### P1 transduction

To combine ME and rRNA deletions with tRNA deletions, we used P1 transduction (Thomason et al., 2007). We grew recipient strains overnight and resuspended in equal volume of MC buffer (100 mM $MgSO_4.7H_2O$ and 5 mM $Cacl_2.2H_2O$) and added 50 µl of phage lysate from the donor strain. After incubating at 37°C (20 mins), we washed the cells twice with 0.1 M citrate buffer (0.06 M citric acid $C_6H_8O_7$ and 0.04 M sodium citrate dihydrate $C_6H_9Na_3O_9$). After the second wash, we resuspended the cells in 1 mL LB with 20 mM sodium citrate and incubated at 37°C (60-90mins). Finally, we pelleted the cells, resuspended in 100 µl of citrate buffer and 5 mM sodium citrate, plated on LB agar with Kanamycin (30 µg/ml) and incubated overnight at 37°C.

#### Removal of kanamycin cassette after generation of mutants

We removed the kanamycin cassette that replaced the deleted gene using plasmid pCP20 (Datsenko and Wanner, 2000). Briefly, we grew transformants (mutant+pCP20) at 30°C, 18 h in LB medium (with ampicillin 100 µg/ml) and streaked them on LB agar plate with ampicillin. We patched individual colonies on LB agar, LB+Amp and LB+Kan and incubated overnight at 37°C. We confirmed the loss of the Kan marker by PCR-amplifying and Sanger sequencing those colonies that had lost antibiotic resistance (indicating loss of the Kan cassette and the Amp-marked plasmid), and grew only on LB agar.

#### Measurement of translation capacity with GFP reporter

We grew the WT and mutants transformed with plasmid pBAD::GFP2 overnight in LB medium with ampicillin for 14-16 hours. We subcultured 1% v/v into M9 with 0.8% glucose and amino acids (with ampicillin) and incubated for around 4-5 hours (mid log phase) before inducing with 0.5 mM IPTG. After induction, we measured GFP and $OD_{600}$ in a microplate reader (Infinite Pro Tecan, Austria). We first normalised GFP readout with OD at the same time point and then took the difference in GFP reading between time zero (immediately after induction) and after one hour to estimate the amount of GFP produced in an hour. We then normalised this readout for each mutant with the WT, to get relative translation capacity of mutants.

#### Measuring tRNA pools with YAMAT-Seq

We extracted total RNA from each growing culture using a TRIzol Max Bacterial RNA isolation kit (Invitrogen™; catalogue number 16096040) as per the manufacturer's protocol. We subjected 10 µg of total RNA per sample to tRNA deacylation by incubating in 100 µl of 20 mM Tris-HCl (pH 9.0) for 40 minutes at 37°C. We then desalted the deacylated RNAs and concentrated by ethanol precipitation. Next, we took 1 µg of each precipitated product and ligated Y-shaped, DNA/RNA hybrid adapters (Eurofins; Shigematsu et al., 2017) to the conserved, exposed 5'-NCCA-3' and 3'-inorganic phosphate-5' ends of uncharged tRNAs using T4 RNA ligase 2 (dsRNA Ligase; New England BioLabs Inc., catalogue number M0239S). We reverse transcribed ligation products to cDNA using SuperScript™ III reverse transcriptase (Invitrogen™; catalogue number 18080093), and amplified cDNA products by eleven rounds of PCR with Phusion® High Fidelity Master Mix with HF Buffer (New England BioLabs Inc., catalogue number M0531S) with sample-specific indices (Illumina). We checked the quality and quantity of each PCR product using an Agilent DNA 7500 kit on a Bioanalyzer and a Florescence Nanodrop, respectively, and combined all samples in equimolar amounts into a single tube. We ran the mixture on a 5% native polyacrylaminde gel, and excised bands between ~200 bp and ~280 bp size, which are expected to be enriched in tRNA-adapter sequences. Finally, we extracted the excised DNA in deionized water overnight, and removed agarose by centrifugation through filter paper. Quality and quantity of the final product was checked on a Fluorescence Nanodrop. The final product was sequenced by the sequencing facility within the Max Planck Institute for Evolutionary Biology, using an Illumina NextSeq 550 Output v2.5 kit (Single-end, 150 bp reads). Raw reads and analysis files are available from NCBI GEO (accession number GSE198606).

**Data analysis**

We used Graphpad Prism (9.0.0) for all statistical analysis (except for YAMAT-Seq) and heatmaps, and DEseq2 (Love et al 2014) and R version 3.6.0 (R Core Team 2021) for the analysis of YAMAT-Seq data as described above.

Datsenko KA, Wanner BL. 2000. One-step inactivation of chromosomal genes in Escherichia coli K-12 using PCR products. *Proc Natl Acad Sci U S A* **97**:6640–6645. doi:10.1073/pnas.120163297

Shigematsu M, Honda S, Loher P, Telonis AG, Rigoutsos I, Kirino Y. 2017. YAMAT-seq: an efficient method for high-throughput sequencing of mature transfer RNAs. *Nucleic Acids Res* **45**:e70. doi:10.1093/nar/gkx005

Thomason LC, Costantino N, Court DL. 2007. E. coli Genome Manipulation by P1 Transduction . *Current Protocols in Molecular Biology* **79**:1–8. doi:10.1002/0471142727.mb0117s79

### SUPPLEMENTARY FIGURES

**Figure S1: Growth curves ($OD_{600}$ vs time) for representative strains in rich and poor media.**

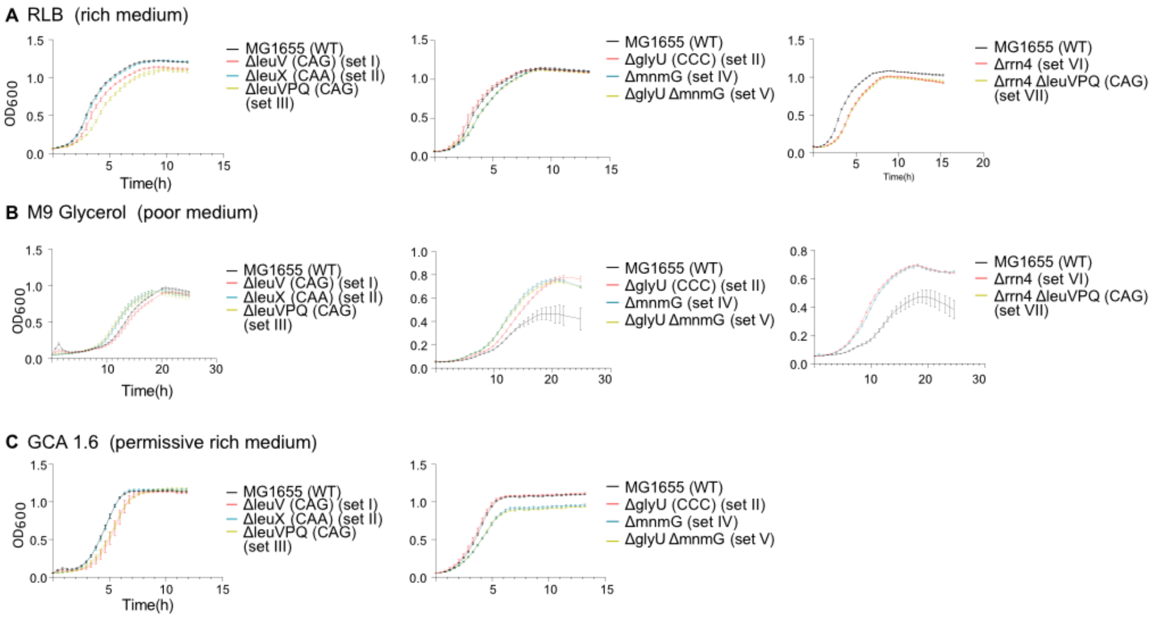


Raw growth curves are shown for one strain from each mutant set (see Table S2 and Fig 1). (A) rich medium (LB, Lysogeny Broth) (B) permissive rich medium (GA1.6, M9 salts + 1.6% glucose + 1.6% cas amino acids) (C) poor medium (M9 glycerol, M9 salts + 0.3% glycerol). Error bars indicate SE across replicates (n = 4 per strain per medium).

**Figure S2: Mean relative growth rate ($R_{rel}$), length of lag phase ($L_{rel}$) and yield ($K_{rel}$) of mutant strains across all tested media.**

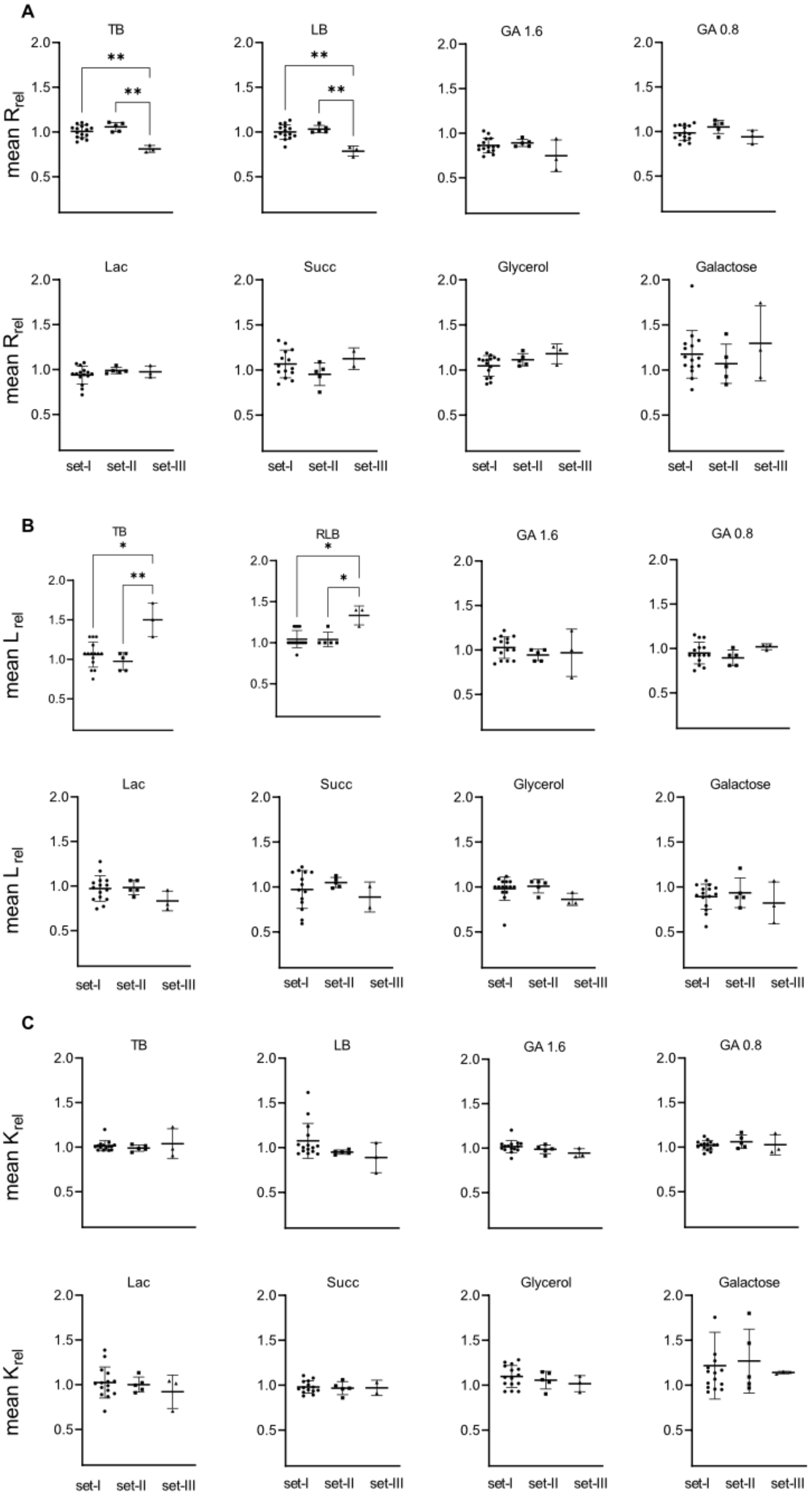

Impact of tRNA deletion on growth parameters (y-axis) for all strains from set I, II and III (on x-axis; see Fig 1, Table S2), shown as the ratio of mutant:WT for each parameter. Each data point is the mean of 4 replicates; the horizontal bar indicates the mean of means of that set. (A) relative growth rate (B) relative length of lag phase (C) relative growth yield. Significantly different pairs (Kruskal–Wallis one-way ANOVA) are showed with asterisks.

**Figure S3: Impact of tRNA deletion on length of the lag phase**.

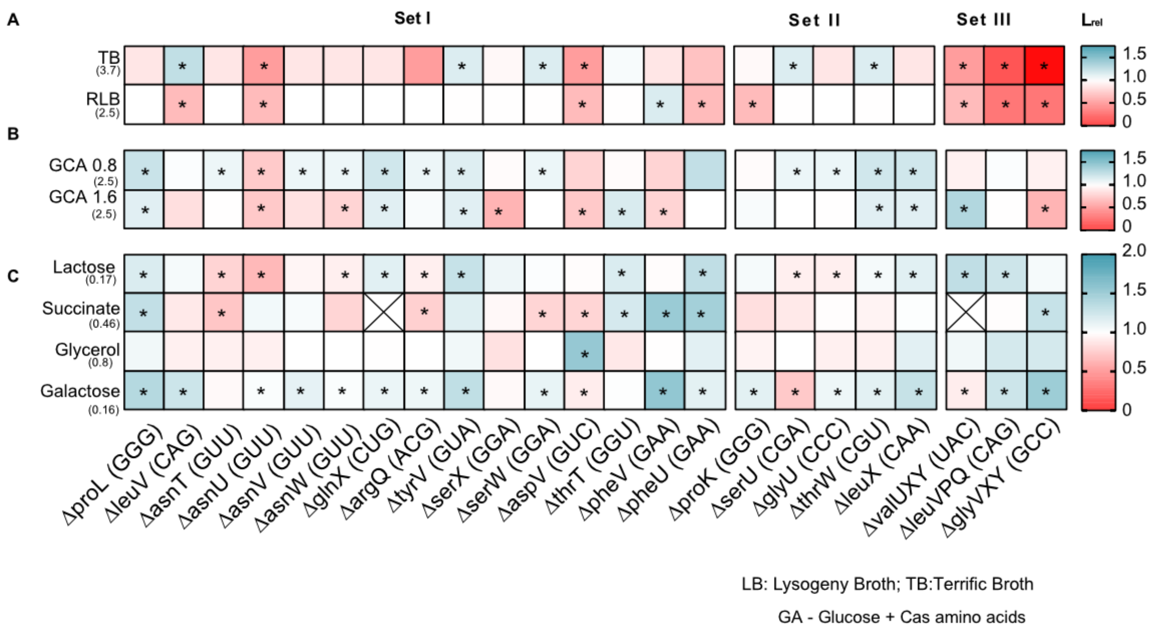


Impact of tRNA gene deletion on the length of lag phase (L), in different media, relative to wild type (WT) ($L_{rel}=L_{deltRNA}/L_{WT}$). The anticodon of each deleted tRNA gene is indicated in parentheses on the x-axis, and strains are categorised into sets as described in Table S2 and Fig 1. Box colours indicate the effect of gene deletion (red: $L_{rel} > 1$, mutant has longer lag phase than WT; blue: $L_{rel} < 1$, mutant has shorter lag phase than WT; n = 3–4 replicates per strain per medium). Asterisks indicate cases where the mutant has a significantly different lag phase than WT (ANOVA with Dunnett's correction for multiple comparisons). The absolute lag length (hours) of WT in each medium is indicated in parentheses on the y-axis. From top to bottom, panels show yield in (A) complex rich media (B) permissive rich media, with indicated concentrations of glucose and cas amino acids (C) poor (M9) minimal media with the indicated carbon source but no cas amino acids.

**Figure S4: Fitness impact of tRNA deletions on growth yield.**

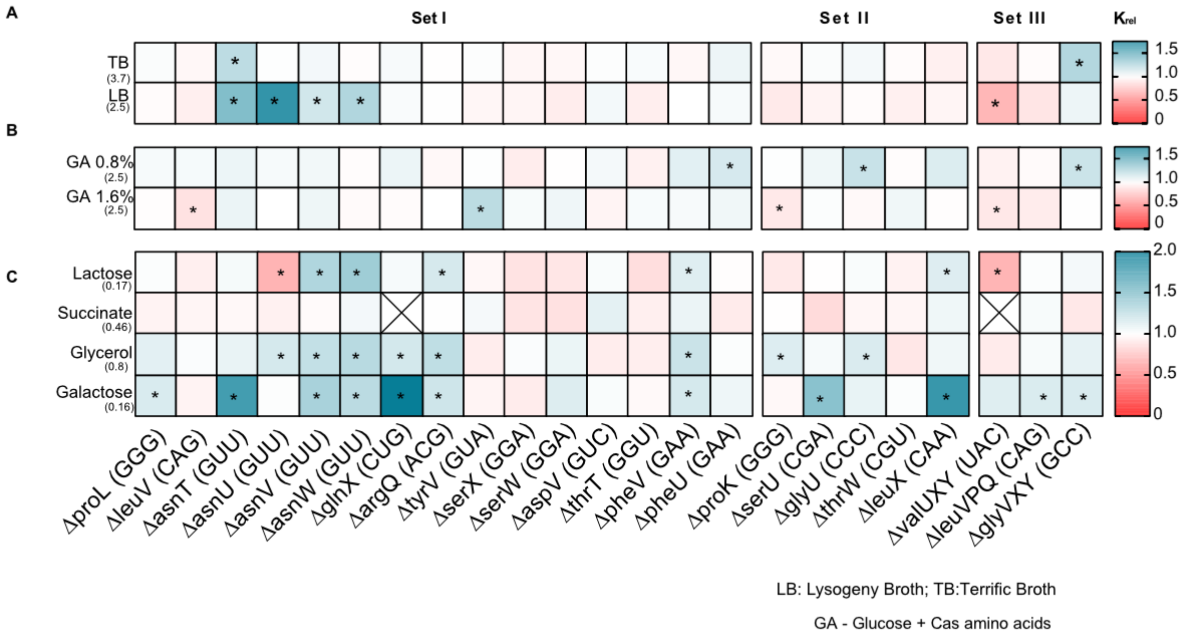


Impact of tRNA gene deletion on growth yield (K), in different media, relative to wild type (WT) ($K_{rel}$=$K_{deltRNA}$/$K_{WT}$). The anticodon of each deleted tRNA gene is indicated in parentheses on the x-axis, and strains are categorised into sets as described in Table S2 and Fig 1. Box colours indicate the effect of gene deletion (red: $K_{rel} > 1$, mutant has higher yield than WT; blue: $K_{rel} < 1$, mutant has lower yield than WT; n = 3–4 replicates per strain per medium). Asterisks indicate cases where the mutant has a significantly different yield than WT (ANOVA with Dunnett's correction for multiple comparisons). The absolute yield (max $OD_{600}$) of WT in each medium is indicated in parentheses on the y-axis. From top to bottom, panels show yield in (A) complex rich media (B) permissive rich media, with indicated concentrations of glucose and cas amino acids (C) poor (M9) minimal media with the indicated carbon source but no cas amino acids.

**Figure S5: Correlation between absolute growth rate of the WT and relative growth rate of mutants.**

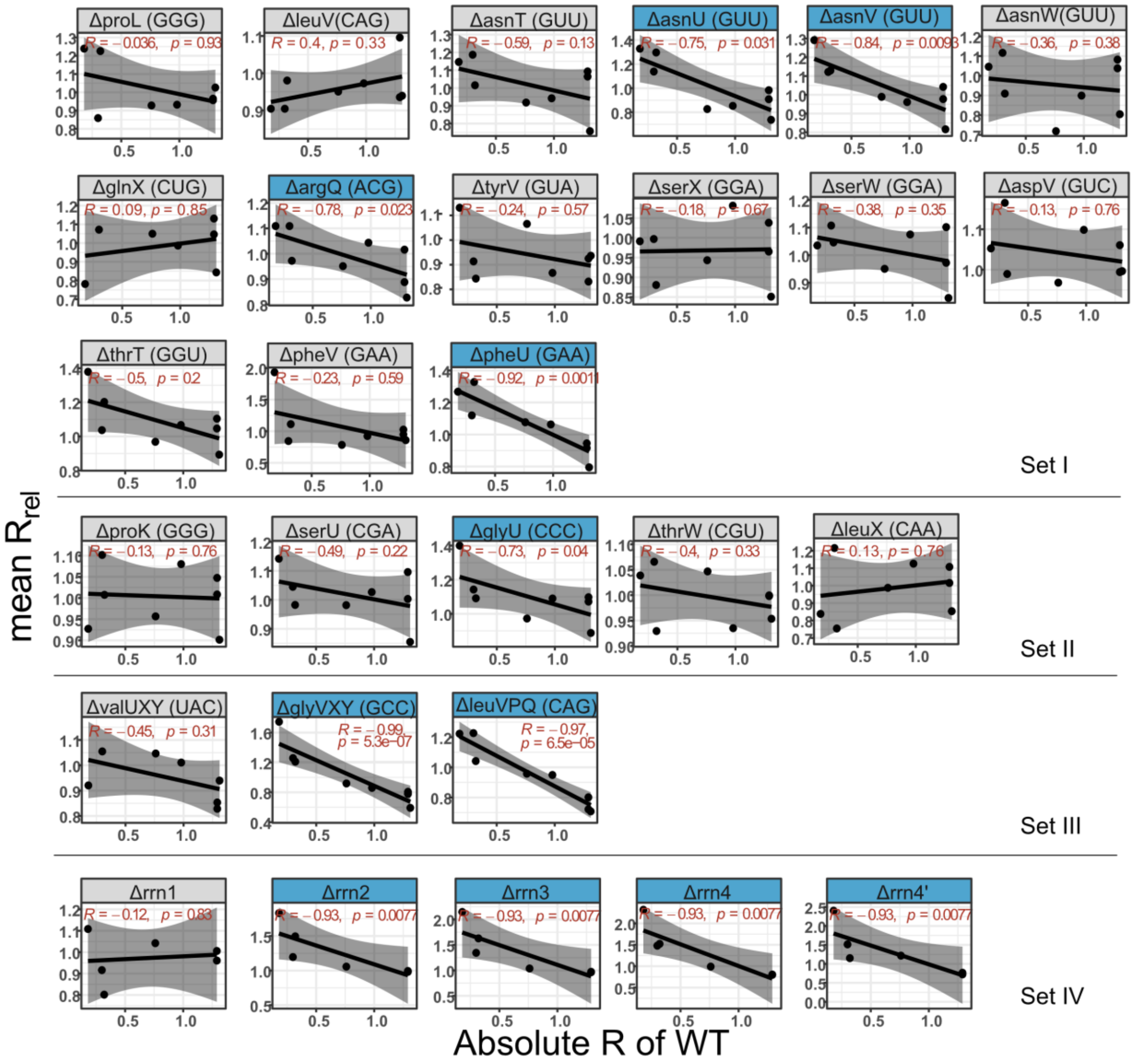


Plots show Spearman's rank correlation between the growth rate impact of each gene deletion across different media ($R_{rel}$) and the respective maximal WT growth rate ($R_{max}$). Strains with statistically significant non-zero slope values are highlighted in blue.

**Figure S6: Relative expression of tRNAs in WT *E. coli* in different media.**

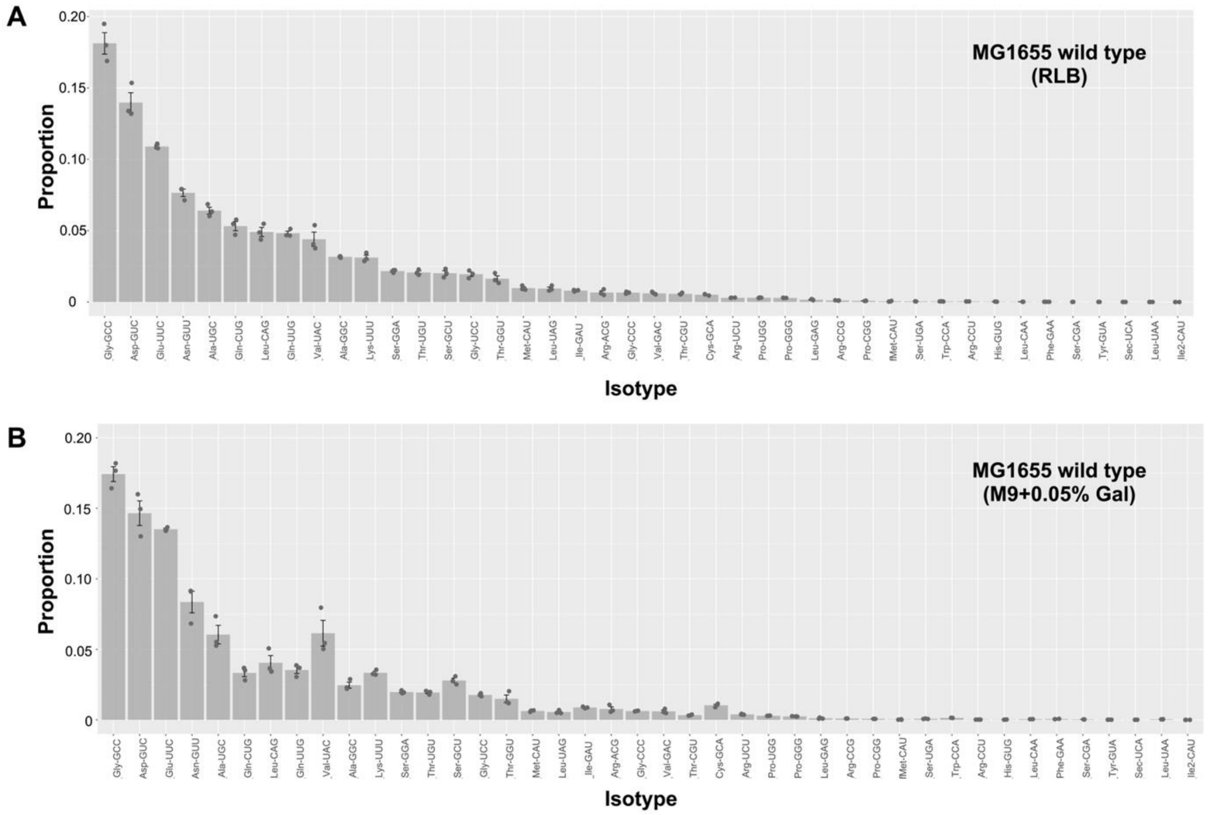


Bar plots show the proportions of 42 tRNA isotypes measured in the mature tRNA pool of *E. coli* K-12 MG1655 (WT) in (A) rich (LB) and (B) poor (M9+0.05% galactose) media. Isotypes are ordered from highest (tRNA-Gly(GCC)) to lowest (tRNA-Ile2(CAU)) in rich medium; *y*-axes scales are the same for both figures. Bars are means of 3 replicates, with individual replicates shown as grey dots. Error bars are +/- 1 standard error.

**Figure S7: Impact of tRNA deletion on levels of tRNA species in rich and poor media.** Relative expression level of tRNAs (mean $log_2$ fold change, n=3 per medium per strain) in mutant compared to the WT. Differential expression of focal tRNAs (indicated on the x-axis) in WT vs. each mutant (tRNA deletion strains indicated on the y-axis), in (A) rich (LB) and (B) poor medium (M9 glycerol). Darker blue colors indicate relatively low expression in mutant, and darker red indicates relatively higher expression. Significant differences are indicated by asterisks. Where expression level is more extreme than indicated in the color key, the fold change value is noted in the box. Expression levels of anticodon carried by tRNA gene deleted in each tRNA deletion strain are indicated by pink borders.

**A**

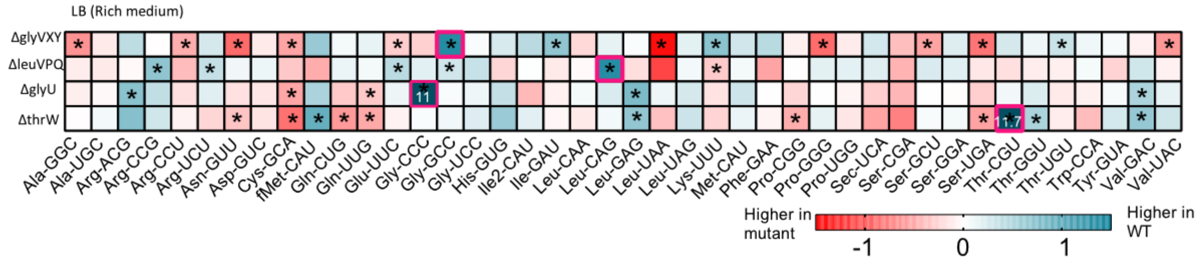


**B**

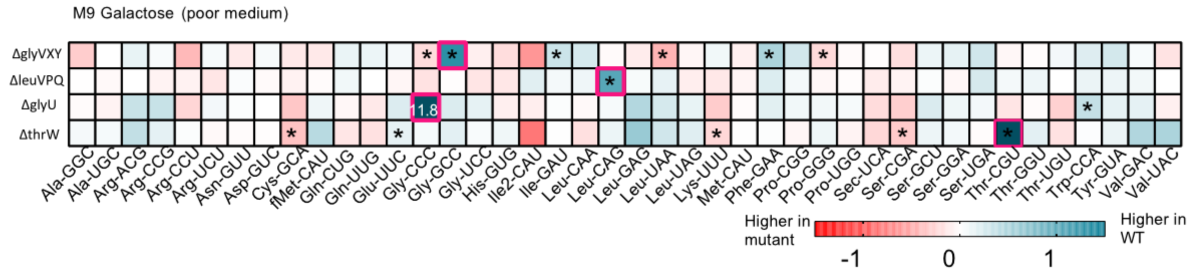

**Figure S8: Impact of tRNA deletions on translation capacity.**

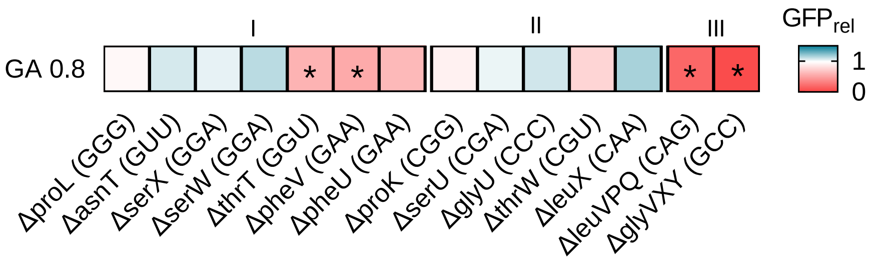


Translation capacity of selected tRNA deletion strains in permissive medium (GA0.8), measured as GFP readout per unit time after induction, relative to WT (n = 2–3 per strain; $GFP_{rel} = GFP_{mutant}/GFP_{WT}$). Red indicates lower production of reporter protein in the deletion strain per unit cell density (i.e. reduced translation capacity), and blue indicates increased translation capacity relative to WT. Significant differences between mutant and WT are indicated by asterisks.
